# Living Neurosheets: Engineering readily deployable neural architectures

**DOI:** 10.64898/2025.11.28.691240

**Authors:** Gaurav Upadhyay, Kimia Kazemi, Seung Hyun Kim, Xiaotian Zhang, Hongbo Yuan, Leo Maslov, Ankit Dhanuka, Zhi Dou, Sehong Kang, M. Taher A. Saif, Hyunjoon Kong, Sergei Maslov, John Beggs, Howard Gritton, Mattia Gazzola

## Abstract

Inspired by the ability of brain structures to support cognitive processing, the need for reliable queryable neural models, and emerging biohybrid applications, we present Living Neurosheets: practical, malleable, modular, and scalable systems for realizing living neural architectures, bio-inspired or beyond evolutionary precedent. By innervating porous sheets, paper-like substrates are endowed with neural function while retaining exceptional versatility. Pre-programmed porosity anchors deposited neurons for reliable spatial control. Cutting, folding, and stacking allow intuitive design, modular composition, and batch fabrication (by the hundreds) of complex neural motifs that preserve geometry and function for months. Reversible integration with tissue-matched optoelectronics enables both sample reuse across hardware and hardware multiplexing across samples, for high-throughput, multimodal investigations. We illustrate these aspects across tissue topologies, gyri-inspired manifolds, and laminar cortical mimics, attaining control over entrainment, synchronization, propagation, suppression, and augmentation dynamics—key features of *in vivo* neural activity. Finally, via cross-instrument portability, we fuse spatial transcriptomics and electrophysiology, linking gene programs to emergent dynamics.

---

Realizing *in vitro* neural architectures with defined spatial cellular compositions, desired processing dynamics, and multimodal interaction capabilities at scale remains a major biotechnology challenge, impairing efforts related to disease modeling, drug screening, biohybrid engineering, and fundamental neuroscience inquiry. Indeed, existing *in vitro* technologies fall short of delivering engineering-level control over tissue design, scalability, and reproducibility—key requirements for efficient prototyping, systematic interrogation, and widespread adoption.

Brain organoids, powerful developmental and disease models (*1, 2*), offer limited structural control and reproducibility (*3, 4*). Bioprinting provides versatility (*5, 6*), however, neural applications remain circumscribed, hampered by material constraints (*5*). In both cases, challenges tied to cell migration and structural integrity persist (*7, 8*), undermining reliable long-term functionality. Further, both technologies suffer from high barriers to entry. Consequently, *in vitro* studies of neural processing continue to rely on 2D cultures (*9–11*), prized for their simplicity yet unable to replicate the hierarchical complexity of native neural tissue.

Multimodal accessibility represents another fundamental bottleneck (*12*). As neural dynamics evolve across molecular, cellular, and network scales through development, plasticity, environmental influence, and disease, their mechanistic dissection requires approaches that not only preserve structure but also enable interrogation across these levels within the same tissue. While standard workflows have been advanced to provide insights into particular aspects of neural tissue dynamics—including connectomics (*13, 14*), proteomics (*15*), genetic profiling (*16*), and electrophysiology (*17*)—these domains remain largely disconnected, typically requiring distinct preparations that preclude same-sample convergent analysis.

Within this context, we introduce Living Neurosheets, malleable biohybrid platforms that offer neural design freedom, streamlined fabrication, and cross-platform interrogation capabilities. Porous sheets—versatile materials with millennia of applications (*18–20*)— are cut to design, innervated, folded, assembled, and optoelectronically interfaced. As intuitive as handling a sheet of paper, this workflow enables architectural control, structural stability (>100 days), and experimental deployment within seconds.

Pre-programmed porosity is exploited to hold deposited cell bodies in place and prevent subsequent migration. This enables robust formation of controlled macro-architectures while allowing neural projections to permeate through the material and fuse into functional 3D networks. As microscopic synaptic connectivity continuously remodels via plasticity, maturation, or disease, this macroscale control decouples structural from functional change (*21*), an essential feature for neuroscience inquiry and computation.

Leveraging these capabilities, we build queryable models of geometry-directed information encoding, synchronization, propagation, suppression, and augmentation—hallmarks of *in vivo* neural activity found in sensory processing, memory, and recall (*22–24*). By expanding the design space through mechanical buckling and folding, we realize motifs inspired by gyri and sulci—structures present in the cortical lamina and associated with higher-order cognitive functions (*25, 26*)—that can be used to decipher the role of curved packings in neural processing. Through modular stacking, we build cortical mimics with cross-layer functional connectivity ablrefere to support propagation and synchronization dynamics.

Lastly, the cross-platform portability of Neurosheets is demonstrated to attain spatiotemporally unified representations that fuse electrophysiology, subcellular resolution spatial transcriptomics, and histology within the same tissue. This approach is employed to track network maturation across molecular, structural, and dynamical scales, revealing coordinated gene expression programs underlying neural recruitment during collective oscillatory dynamics that emerge in mature culture systems.

Thus, by combining programmability, scalability, and cross-platform versatility, Living Neurosheets establish a versatile framework for engineering neural tissues of durable architecture, consistent functionality, and multimodal accessibility, allowing to connect tissue geometry, molecular state, cellular organization, and network activity, with implications in neuroscience, medicine, computing, and engineering.

## Bringing sheets to life

Figure 1 presents an overview of our neurosheet fabrication strategy. We begin by selecting biocompatible materials with pore sizes tailored to simultaneously confine cell somas, enable efficient hydrogel infusion, and support neurite extension (both laterally and orthogonally through the material) to achieve three-dimensional connectivity inspired by the radial interfacing of the allocortex and neocortex (*27, 28*). With simplicity and broad accessibility in mind, we consider two cost-effective and widely available solutions: fiber-reinforced cellulose acetate filters and track-etched polycarbonate filters, with pore diameters ranging from 0.8 to 20 *µm* and thicknesses ranging from 32 to 135 *µm*. While optimal porosity and thickness depend on the application, we report in Figs. 1–4 results based on 100 *µm*-thick cellulose acetate filters of 5 *µm* pore size, selected for their versatility across experimental conditions. An application using 32 *µm*-thick polycarbonate filters with 20 *µm* pores is highlighted in Fig. 5 as an alternative.

**Figure 1.**
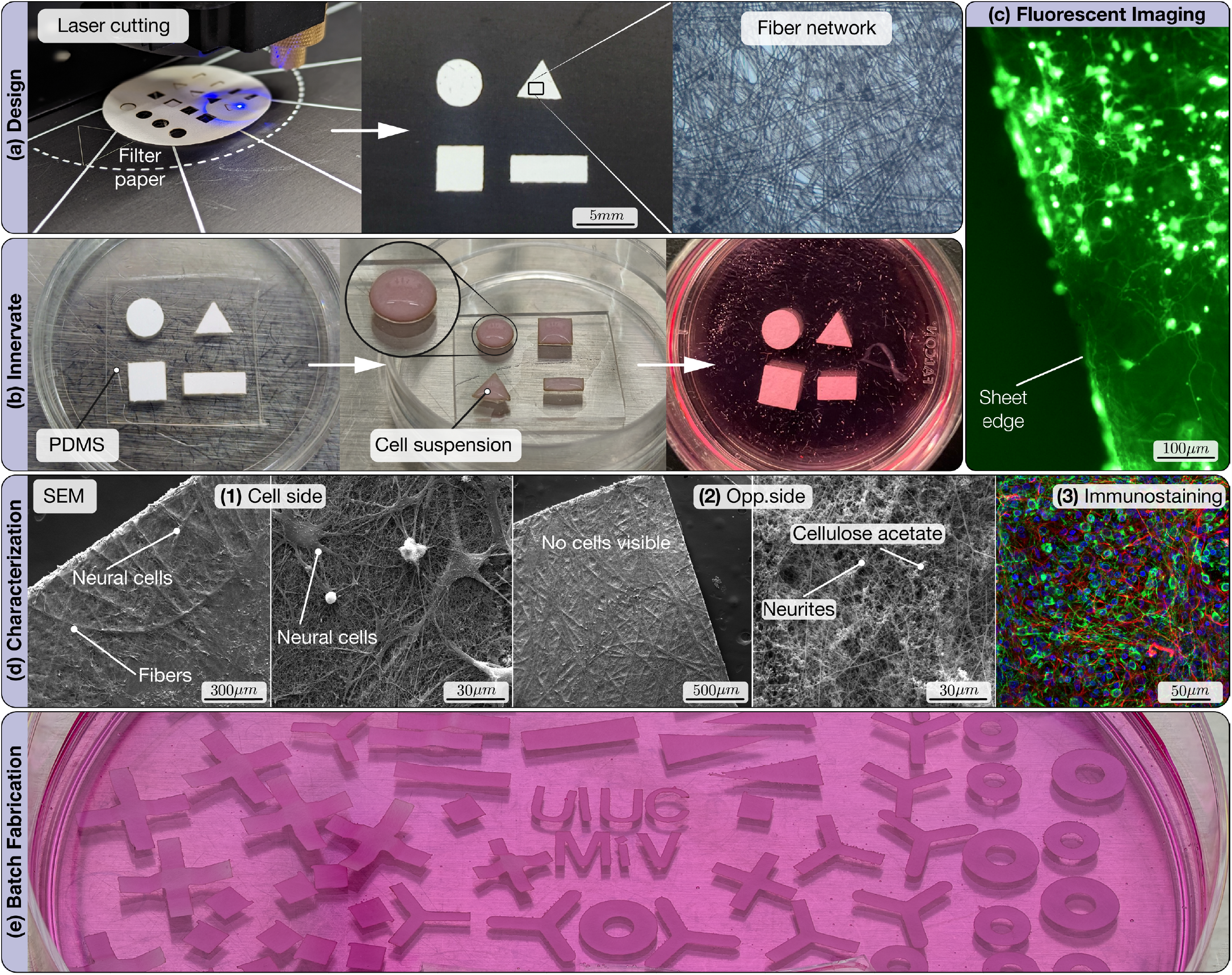
Living Neurosheets. **(a)** Sheets are patterned into formal designs using a laser cutter, then plasma-treated to enhance hydrophilicity. Here, we utilize cellulose acetate membrane filters with reinforcing fibrous architecture, which is visualized using optical microscopy. **(b)** Before cell seeding, samples are placed on anti-cell-adherent coated PDMS. Hydrogel is infused and allowed to gel prior to adding the cell mix, followed by maturation in appropriate media. **(c)** Fluorescence microscopy with ChR2-HBG3 Hb9-GFP mESCs-derived motoneurons enables live imaging of network maturation, with both cellular network and sheet visible. **(d)** (1−2) SEM imaging of both sides of neurosheets confirms their 3D architecture with planar confinement of cell somata on one side, while neurites are present throughout the sheet. (3) Immunofluorescence of primary hippocampal cultures shows homogenous distribution of neurons (green), astrocytes (magenta), and nuclei (blue). **(e)** Batch fabrication across is achieved through a single workflow based on intuitive manipulations such as cutting, folding, and stacking (see also SI. 7)

After selecting the material of choice, corresponding sheets are cut to the desired design using either a benchtop laser cutter or scissors (Fig. 1a). The resulting cutouts undergo sterilization, oxygen plasma treatment to enhance hydrophilicity, and coating with a positively charged polymer to facilitate hydrogel integration and promote cell adhesion. These treated constructs are then placed onto a hydrophobic polydimethylsiloxane (PDMS) substrate to establish a wettability contrast (Fig. 1b). This contrast enables precise confinement of liquids within the seeding area—causing droplets to bead up—thus ensuring accurate control over the deposited hydrogel volume and cell density.

Next, Collagen–Matrigel hydrogel is infused to provide defined mechanical and biochemical cues that support the maturation of neuronal networks (*29*). The resulting sheet–hydrogel constructs, which will serve as tissue culture substrates, can be batch fabricated (Fig. 1e), characterized, and stored—enhancing robustness and scalability. Modularity also permits tapping into the existing infrastructure for sheet functionalization, from microfluidic integration (*19*) to flexible electronics patterning (*20*).

Neurons to be seeded are concurrently prepared following standard protocols (Methods 1.3). To demonstrate the robustness and versatility of our approach, we employ diverse neuronal sources throughout this work, including opsin-expressing and wild-type stem cell–derived motoneurons, as well as primary cortical and hippocampal neurons freshly isolated from rat tissue or commercially procured. Once prepared, neurons are seeded directly onto the hydrogel–sheet constructs in suspension, a straightforward procedure that ensures high cell viability (Fig. S1), uniform cell distribution (Fig. 1d-3), and control over cell density (Fig. S1).

As networks mature into functional biohybrid neural tissues, we confirm via fluorescence imaging (Fig. 1c, Fig. S1, S2) and scanning electron microscopy (Fig. 1c) that neuronal somata are retained at the gel–sheet interface. This localization results from the interplay between porosity and hydrogel mechanics. Confinement of somata enables both geometric control and direct optical and physical access, facilitating high-resolution imaging, patch-clamp and multielectrode-array electrophysiology, optical/chemical stimulation, and molecular extraction (RNA/DNA)—rendering the system highly inspectable.

Importantly, neurites can infiltrate the soft matrix embedded within the porous sheet volume, forming 3D networks. Figure 1d-2 shows neurites extending from the plated soma side through the sheet, reaching the opposite side, which remains inaccessible to cell bodies. Neurite projections can be modulated via hydrogel composition and sheet porosity to form surface- or volume-dominant networks. This provides control over inter- and intra-layer connectivity (Fig. S1, SI. 3), supporting (as we will see) the development of advanced radial multilayer constructs (Fig. 5).

## Optoelectronic interfacing

To demonstrate optoelectronic reversible interfacing, neural recording, stimulation, and imaging, we first consider square-shaped neurosheets.

Bi-directional communication between biological and electronic systems is established by deploying neurosheets on Mind in Vitro (MiV) electrophysiology platforms (*17, 30*). These open-source systems are selected for their accuracy, robustness, affordability, modularity, and customizability. They enable a complete signal chain, stretching from neural cultures and data acquisition to cloud-based advanced analysis and storage—providing an ideal foundation for scalable and reproducible experiments.

In Fig. 2, mature neurosheets innervated with opsin-expressing, mouse stem cell-derived motoneurons, are transferred from the culture dish onto a custom 512-channel microelectrode array (MEA) MiV system (Fig. 2a-1, Fig. S3). Sheets are positioned with the cell-seeded side facing downward to ensure contact between somas and recording electrodes. To secure the tissue and prevent shifting during recording, gentle pressure is applied using a Kimwipe moistened with culture medium (Fig. 2a-2). After filling the MEA with medium to maintain electrical grounding and support tissue viability, experiments can commence immediately.

**Figure 2.**
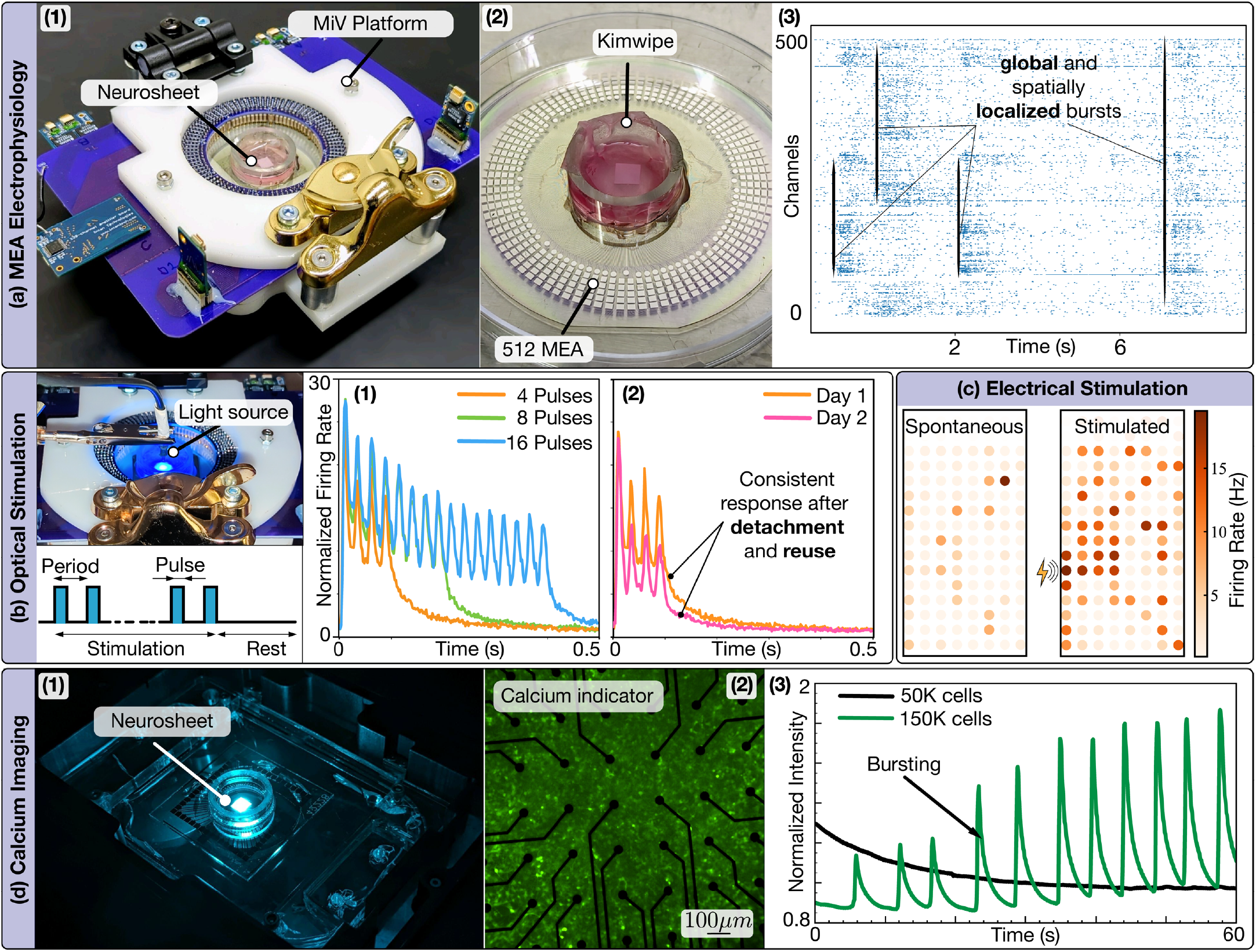
Functional Characterization. **(a)** Electrophysiology pipeline: (1) Open-source Mind in Vitro systems coupled with square neurosheet. (2) Tissue is placed with the cell-seeded side facing the microelectrode array and gently pushed using a Kimwipe. (3) Raster plot showing spontaneous activity with local and global avalanches. **(b)** Optical encoding of information. Encoding is performed using a top-mounted laser with a pulsed stimulation protocol. (1) Normalized Peri-Stimulus Time Histograms (PSTH) reveal distinct peaks corresponding to pulse number. (2) A 4-pulse protocol elicits near-identical responses after reattaching the tissue on day 2. **(c)** Localized electrical stimulation elicits tissue-wide, distance-dependent response. **(d)** Calcium imaging: (1) MEA with neurosheet can be directly mounted on the microscope for imaging. (2) Calcium indicator labels cells green and remains visible alongside the MEA. (3) Primary cortical neurons exhibit density-dependent dynamics, with lower densities showing mostly asynchronous activity and higher densities producing periodic synchronized responses.

The spontaneous neural activity recorded from a representative sample is visualized in Fig. 2a-3 as a raster plot, illustrating spike events across electrodes over time. The data exhibit a rich repertoire of spatiotemporal dynamics, with activity self-organizing into cell-specific spikes as well as subnetwork and pan-network bursts—consistent with observations in organotypic brain slices (*17, 31, 32*). We empirically observe that the spatial extent of activity propagation is modulated by the pore size, with smaller pores promoting localization and larger pores conducive to distal propagation (SI. 3, SI Video 1). Neurosheets remain viable for up to 100 days, preserving tissue architecture and enabling longitudinal tracking of dynamic changes over months and across samples (SI. 1, Fig. S6), to study maturation/aging.

Next, we demonstrate the neurosheets’ ability to encode multimodal inputs, delivered electrically or optically. The latter is enabled by the use of opsin-expressing neurons that respond to blue (473nm wavelength) light. Optical inputs are provided by a laser source positioned on top of the MEA, delivering trains of 4, 8, or 16 pulses (10ms-long, 40Hz), interspersed by 3-second intervals (Fig. 2b). Evoked neural responses are recorded and visualized in Fig. 2b-1, using global peristimulus time histograms (PSTH). As shown, pan-network activity results in well-defined peaks that track the input—a key feature to embed information and harness dynamics for sensory processing (*33, 34*).

Unlike conventional tissue-engineered models that require permanent MEA bonding, neurosheets can be attached/detached within seconds. This is showed in Fig. 2b-2, where a tissue is tested, removed from the MEA, stored, and repositioned the next day, yielding near-identical responses and confirming robust re-engagement. This reversible integration with electrophysiology systems is a distinctive advantage of neurosheets, transforming static interfaces into reusable, high-throughput platforms. Tissue portability indeed enables longitudinal studies across many samples with a single MEA device. It also supports cross-platform workflows (SI. 2), facilitating convergent analyses that combine electrophysiology, imaging, or omics perspectives, as discussed later.

Neurosheets’ versatility is further demonstrated through electrical and chemical stimulation as well as imaging. In Fig. 2c, we examine the response of a tissue to an electrically localized stimulus. Firing rate heatmaps indicate a post-stimulus, spatially distributed increment of neural activity. The response global maximum is found at the stimulation site, demonstrating network-wide entrainment to locally embedded information. Similarly, chemical stimulation is shown to modulate global dynamics (Fig. S6). Complementing MEAs’ high temporal resolution, neurosheets’ optical accessibility enables calcium imaging for single-cell spatial detail. For illustration, multiple samples on a microscope-integrated MEA (Fig. 2d-1,2) are imaged to investigate burst dynamics versus cellular density. As shown in Fig. 2d-3, higher-density cultures exhibit regular synchronization, as opposed to lower-density ones, consistent with prior studies (*35*). Density-dependent control can then be exploited for tailoring neurosheets to specific functional tasks.

## Planar tissue topologies

Next, we manipulate tissue topology and geometry for neural dynamics control. The ‘directed emergence’ of (intractable) microscopic connectivity via macroscopic boundary conditions is a key step toward spatially programmable neural architectures of prescribed function. We illustrate this concept through examples of information transport, suppression, segregation, and augmentation.

To impart dynamics of transport, directionality, and integration, we move beyond square tissues and adopt configurations with branching and merging. We begin with a ring motif—characterized by a periodic boundary condition—that allows inputs to diverge, converge, and self-interact (Fig. 3). Ring-like motifs are common across neural systems, from the fly central complex, encoding head direction, to cnidarian nerve nets around the oral or bell margins of jellyfish, coordinating feeding and locomotion (*36, 37*). Ring topologies are cut, treated, and innervated with either wild-type or ChR2-transfected motoneurons (Fig. 3a-1). Guided by observations in rectangular strip systems (SI. 4), the ring’s centerline diameter is set to 7 mm—large enough to prevent spontaneous global synchronization and enable long-range activity propagation, yet small enough to allow dynamics to spatially interact and to limit dissipation. Figure 3a-1 shows an immunostained sample after 14 days of maturation, where neurons and astrocytes are distributed homogeneously—a prerequisite for maintaining continuity of information flow. Finally, to enable precise electrophysiology, ring-shaped, high-density MEAs (Fig. 3a-2) are custom fabricated and tissue-matched.

**Figure 3.**
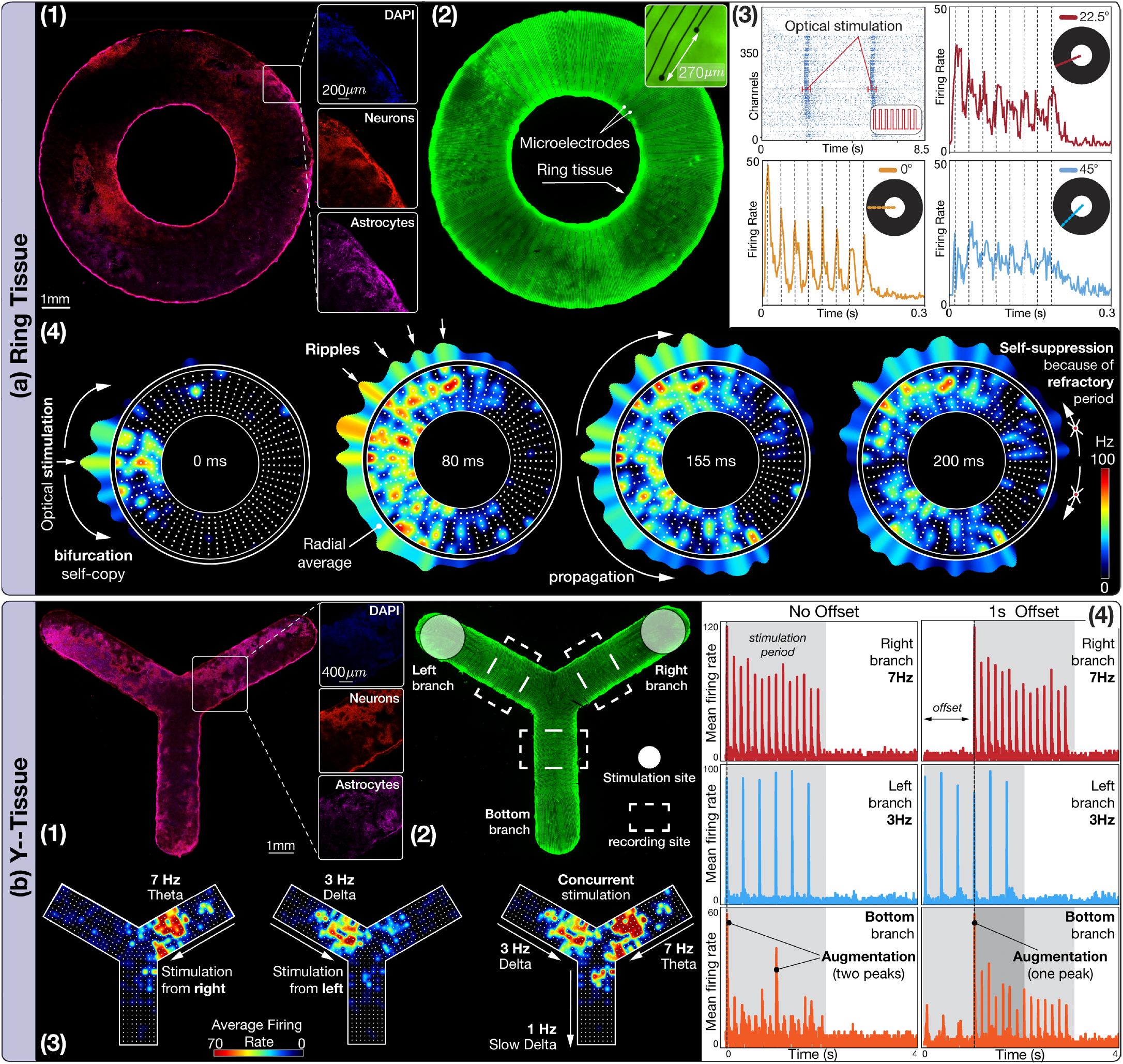
Planar tissue topologies. **(a) Ring Tissue.** (1) Immunostaining shows nuclei (blue), neurons (red), and astrocytes (purple). See SI Video 2. (2) Ring tissue (GFP neurons, green) coupled with tissue-matched, high-density microelectrode array. (3) Pattern encoding and propagation. Raster plot shows tissue-wide propagation upon localized optical stimulation (in red). PSTH plots visualize the mean firing rate at various locations along the circumference of the ring, post stimulation. Dashed lines correspond to peaks encoded at the stimulus onset location. (4) Averaged heatmaps of the mean firing rate after stimulation. Input (from the left) is encoded by the tissue as ripples, which bifurcate and propagate before meeting and canceling out at the circumferentially opposite end (SI Video 4). Similar dynamics are observed across samples and stimulus locations (Fig. S4) **(b) Y-Tissue**. (1) Immunostaining shows nuclei (DAPI), neurons (red), and astrocytes (purple). See SI Video 5. (2) Y-tissue (GFP neurons, green) coupled with tissue-matched MEA. Stimulation sites and locations for estimating peri-stimulus histogram (PSTH) plots are highlighted. (3) Heatmaps of the mean firing rate activity after stimulation. Light pulses at 7 Hz (theta) and 3 Hz (delta) are encoded in the right and left branches. Signal augmentation happens at a 1 Hz (slow delta) in the bottom branch (SI Video 8). (4) PSTH plots show location-specific mean firing rates. The left column shows branch encoding when input streams are provided without offset. The right column corresponds to a 1-second offset between left and right branch stimulation. Similar behavior is observed across multiple samples (Fig. S9). For segregation dynamics in Y-tissues, see SI Video 7.

We hypothesize that toroidal tissues—which can effectively be thought of as periodic, axisymmetric branches—will support bifurcation dynamics, parallel transport, and two-stream interaction. Such systems thus represent minimal models for investigating mechanisms of signal copy, redundancy, propagation, or refractory period localization, all mechanisms biologically and computationally relevant (*38, 39*). This intuition is supported by neural recordings in maturing rings (SI Video 3), where spontaneous avalanche events are found to bifurcate into symmetric clockwise and counterclockwise propagating fronts.

To assert command over these emergent dynamics, we externally induce local avalanches through spatially targeted optical stimulation (Fig. 3a-3,4), using the pulsed protocol of Fig. 2. Similar to the square neurosheet, the ring tissue reliably encodes each pulse, producing distinct peaks, visible in the PSTH plots at the stimulus site (Fig. 3a-3). Independent of the stimulus location (tested throughout the ring, Fig. S4b,c), locally encoded events immediately self-copy and bifurcate into counterpropagating wavefronts (Fig. 3a-3), reflecting the inherent symmetry of the ring and closely resembling dynamics induced by spontaneous avalanches. Notably, optically-initiated traveling waves preserve the input’s temporal structure in the form of radially aligned ripples with fixed temporal offsets (Fig. 3a-3,4), consistent with large-scale brain activity coordination *in vivo* (*40–42*). As waves propagate, the ripples progressively distort due to dissipation, network recurrence, and geometric dispersion arising from the mismatch between inner and outer arc lengths. By reasoning that larger input energy might allow wavefronts to travel farther, we increase the number of pulses from 4 to 8. We observe that under the 4-pulse condition, ripple amplitudes vanish before reaching the ring’s opposite site, preventing the interaction between clockwise and counterclockwise fronts (Fig. S4b). In contrast, the 8-pulse stimulation of Fig. 3a-4 allows bifurcated waves to meet at the opposite end, albeit attenuated. Upon collision, activities from both directions cancel out due to the refractory zones (regions in recovery after spiking) trailing each ripple, realizing a mechanism of self-suppression. These rich dynamics render the ring feedforward design a useful precursor model to study other relevant neural motifs, including feedforward inhibition and excitatory-inhibitory recurrent architectures (*43, 44*).

Observations of active and refractory dynamics in the ring topology motivate our next motif: the Y-junction architecture of Fig. 3b. While the ring serves as a blueprint for exploring same-stimulus bifurcation, self-interaction, and suppression, the Y-junction allows us to study the interaction of independently-generated inputs, originating from distinct neural populations: a minimalistic model of sensory systems where rhythms emerging across different brain regions are fused, integrated, or segregated.

Following the same workflow as for the ring topology, Y-constructs are cut, treated, and innervated with either wild-type or ChR2-transfected motoneurons. Drawing on insights from both strip and ring preparations, the lengths of the three Y-branches are adjusted so that neural activity evoked at the distal ends reliably propagates to the central junction (SI 6). Again, immunostaining confirms the homogeneous distribution of neural cells throughout the tissue, and topology-matched MEAs are fabricated to enable precise electrophysiology (Fig. 3b-1,2).

After confirming the spontaneous emergence of localized avalanches and propagation fronts (SI Video 6), we optically drive rhythmic neural activity at the top branches, to examine stream fusion at the junction and resulting output at the downstream branch. For this demonstration, we applied 3 Hz (delta) and 7 Hz (theta) stimulations for 2 seconds at each top branch, motivated by the well-established co-occurrence and cross-frequency coupling of delta–theta rhythms during sensory processing (*45, 46*). As in previous motifs, each input elicits robust, frequency-locked neural firing near the stimulation site, with 2-second PSTH plots showing 6 peaks for 3 Hz and 14 peaks for 7 Hz (Fig. 3b-3,4).

When stimulation at the top branches is simultaneously initiated, both theta and delta activities are observed to travel through the junction and co-exist in the downstream branch. The PSTH plot collected at the output branch indeed presents distinct peaks, phase-locked to each frequency of stimulation. Notably though, two time stamps—exactly at 0 s and 1 s—display markedly amplified responses. These events align precisely with the coincident arrival of both delta and theta fronts at the Y-junction, revealing a geometry-imposed, timing-dependent amplification mechanism. This is confirmed by offsetting the two inputs by 1 s, which indeed produces only a single augmented peak. These dynamics thus implement an autonomous coincidence detection mechanism, where enhanced activation is achieved only when two signals arrive within a narrow temporal window (*47*). Coincidence detection is a prominent feature of auditory sensory processing and sound localization (*48*). Just as important, outcompetition by early-arrival signal is a prominent feature of distractor suppression in visual fields (*49, 50*). The Y-design thus provides a basis for selective attention models to study competition between receptive fields for attentional augmentation.

These examples demonstrate that neurosheets can be prototyped to probe principles of rhythmic emergence, theorized as part of neural computation strategies, but difficult to test *in vivo*.

## Tissue manifolds

Having established planar spatial programmability, we expand the design space through folding and buckling to create functionally innervated 3D manifolds—compliant neural structures unattainable with current approaches. We are motivated by biological systems that exploit curvature and folds—most notably cortical gyri and sulci—to achieve dense packing and support rich connectivity. Moreover, from a biohybrid robotic perspective (*51, 52*), integrating ‘nervous systems’ into deformable, motile platforms demands neural architectures that dynamically conform while maintaining functional integrity. To address these challenges, drawing from origami and kirigami reconfigurable topologies, we engineer planar neural tissues that transform into curved geometries, enabling out-of-plane signal routing while preserving robust propagation.

We begin by inducing buckling of a planar paper strip via compressive forces applied from both ends, resulting in controlled mechanical snapping at the sheet’s center to form an arch bridge configuration (Fig. 4a-1). Once buckled, the sheet is infused with hydrogel and innervated with neurons—either wild-type or ChR2-transfected—which distribute uniformly throughout the tissue (Fig. 4a-2). After 14 days of maturation, the construct is positioned on MEAs aligned with the arch’s legs (Fig. 4a-1) for electrophysiological assessment of neural functionality. This is indeed confirmed in Fig. 4a-3, where spontaneous avalanches are detected at both legs, with a slight delay suggesting tissue-wide neural activity propagation from Leg-1 to Leg-2.

**Figure 4.**
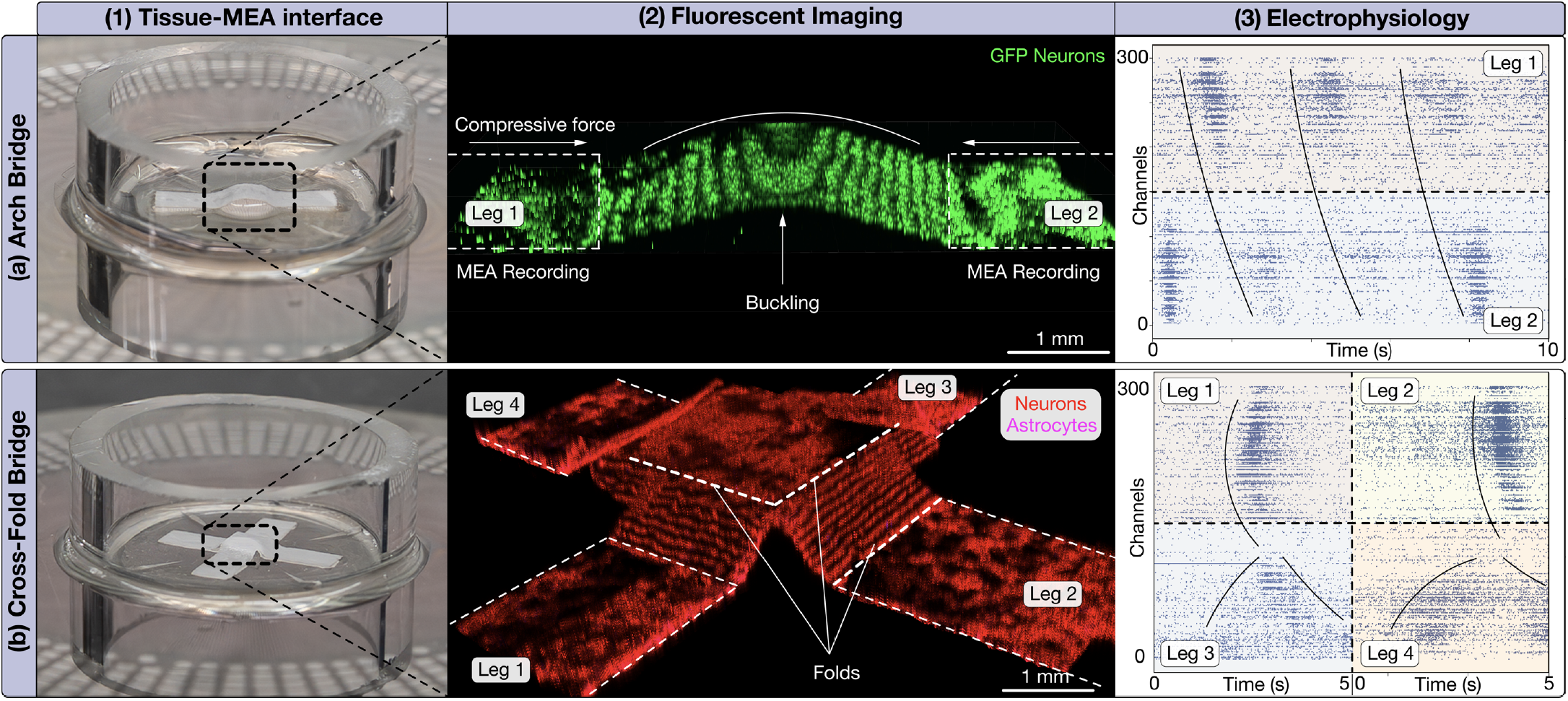
Tissue manifolds. **(a)** Arch bridge configuration realized by applying equal force from both ends of a rectangular strip, (1) buckling the sheet from the middle and creating a smooth curvature. (2) The innervated tissue is imaged via fluorescent imaging. (3) When coupled with MEAs, both legs reveal spontaneous avalanches, which appear with a slight delay, signaling tissue-wide communication. **(b)** Cross-Fold bridge configuration realized by (1) folding the legs of a cross-shaped planar sheet, creating sharp creases. (2) The innervated tissue is immunostained for neurons and astrocytes and imaged via fluorescent imaging. (3) Coupling with MEAs is performed with two opposite legs (Legs 1 and 3, Legs 2 and 4) at a time. All four legs exhibit spontaneous avalanches, appearing with slight delays, signaling tissue-wide communication. The Z-scale of b2 is slightly expanded to better visualize the structure after fluorescent imaging compression (SI 2.9).

**Figure 5.**
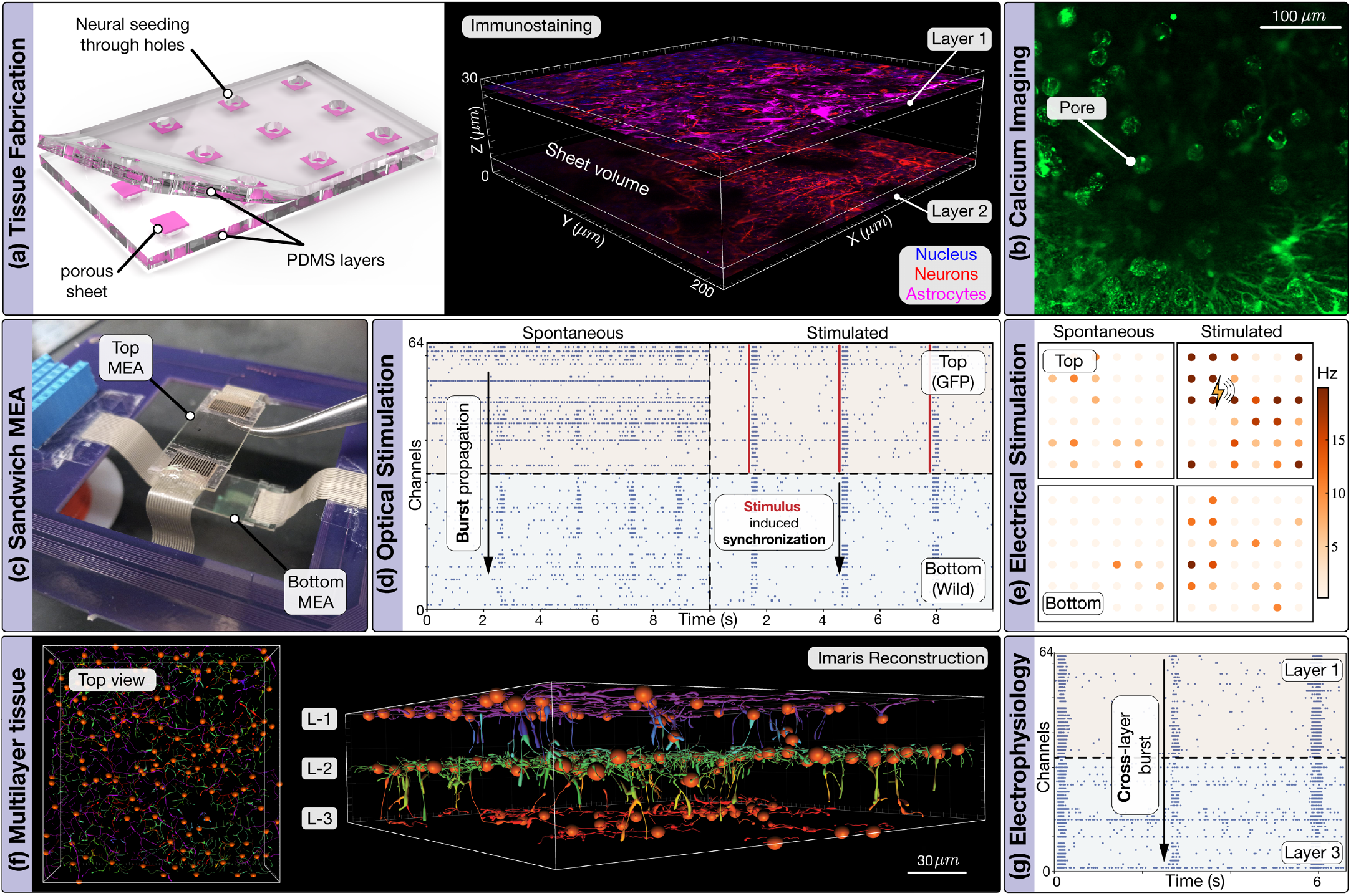
Laminar tissues. **(a)** Tissues are made by sandwiching porous sheets between through-punched PDMS layers. Immunostaining shows that cells remain on each sheet side with neurites penetrating the pores, see SI Video 9. **(b)** Calcium imaging shows neural activity within pores. **(c)** The sandwich MEA uses two MEAs on one PCB to record simultaneously from top and bottom layers. **(d)** Tissues with ChR2-GFP motoneurons on one side and wild-type on the other display spontaneous avalanches with both inter- and intra-layer propagation. Optical stimulation phase-locks avalanches to the stimulus period. **(e)** Electrical stimulation via sandwich MEA induces a localized perturbation that propagates across sides. **(f)** Imaris reconstruction of confocal imaging data of trilayer constructs showing cells’ somas in respective planes with neurites spanning the entire tissue, see SI Video 11. **(g)** Electrophysiology on three-layer constructs using the sandwich MEA shows cross-layer burst propagation. See SI Videos 12-14 for full characterization and Fig. S11 for consistency across cell types.

Complementarily, neural 3D manifolds can also be realized through folding. As an example, in Fig. 4b-1, a four-legged cross-bridge architecture is created by folding a planar cross-shaped motif. Following creasing, the construct is infused with hydrogel and uniformly seeded with neural cells (Fig. 4b-2). After maturation, the tissue is transferred onto MEAs to record neural activity from the legs. As shown in Fig. 4b-3, spontaneous avalanche events occur across all four legs, with inter-leg delays indicative of long-range, cross-talk interactions within the folded geometry. As with the arch bridge, full spatiotemporal dynamics quantification across the manifold can only be obtained via out-of-plane MEAs, which remains an area of active research.

Together, these architectures demonstrate that planar neural constructs can be mechanically configured into curved and folded manifolds of varying topology, preserving functional activity. Indeed, both buckled and folded systems are found to support spontaneous, network-wide dynamics, underscoring the persistence of long-range coordination across the manifold.

## Laminar tissues

Laminar circuit organization is a defining feature of many sensory and associative structures—including the cerebral cortex, retina, or hippocampal formation—where microcircuits are stacked into layers, enabling the hierarchical transformation and integration of signals. Motivated by these biological exemplars, we engineer neurosheet stacks that recapitulate laminar motifs in a modular and durable manner, paving the way to examining the emergence, propagation, and interaction of neural rhythms across discrete, but connected, layers.

We begin by establishing a bilayer motif in which neural cells are seeded on both sides of a hydrogel-infused polycarbonate sheet. Owing to their minimal thickness (32 *µ*m), polycarbonate sheets support modular stacking without exceeding diffusion limits. Further, their cylindrical micro-pores guide neurite extension perpendicular to the sheet, increasing the likelihood of interlayer connectivity. For robust and scalable fabrication, up to 12 membranes are secured between two punched PDMS layers (Fig. 5a), enabling parallel processing of multiple samples in isolated compartments. Cell seeding occurs through the PDMS openings, each exposing a 5mm-wide, circular region of the underlying sheet. This design prevents lateral cell leakage, allows batch handling without direct contact with seeded surfaces, and preserves tissue integrity throughout maturation.

Seeded tissues are allowed to mature, with porosity confining cell somas to the opposite sides of the sheet while permitting neurites extension through the membrane volume. Bilayers are innervated using wild-type, ChR2-transfected motoneurons, or a mixed configuration with wild-type cells on one side and optogenetic neurons on the other. By day 14, immunostaining reveals uniform distributions of neural cells on both sides (Fig. 5a). Calcium imaging further shows neural activity on each side and communication through the sheet’s pores (Fig. 5b, SI Video 10).

To perform electrophysiology characterizations of functional connectivity in laminar constructs, we devise a multi-layer MEA strategy. This relies on the use of two planar MEAs that ‘sandwich’ the tissue, providing top and bottom interfacing. This approach underscores how neurosheets support bidirectional development of both novel tissue architectures and interfaces, thereby systematically advancing neural systems engineering.

Once interfaced, mature constructs exhibit spontaneous avalanches propagating across the two cellular layers, indicating functional inter-population connectivity (Fig. 5d). To assess responsiveness to external stimulation, we employ heterogeneous bilayers equipped with optogenetic (top side) and wild-type neurons (bottom). By selectively light-activating the top layer, optically evoked avalanches are observed to reliably traverse the construct and entrain the input at the bottom layer (Fig. 5d). Similar results are obtained with electrical stimulation (Fig. 5e).

With bilayer modules established, we construct higher-order architectures by stacking multiple bilayers separated by thin hydrogel interfaces. Using primary rat or mouse cortical neurons, we thus form ‘cortical stacks’, imaged and tested in Fig. 5f,g. Specifically, two bilayer units are combined to form a trilayer, reflecting allocortical structures in reptiles and mammals (*27, 28*). Upon maturation, calcium imaging confirms activity across all three layers (SI Videos 12,13), while sandwich MEA recordings reveal spontaneous avalanche propagation through the stack, demonstrating inter-layer coupling. Notably, these laminar architectures preserve their macrostructure for the entire sample’s lifetime (months), overcoming current technological limitations.

## Spatiotemporally fused multiscale and multimodal representation of neural function

We conclude by leveraging the cross-platform portability of Neurosheets to integrate high-resolution electrophysiology, subcellular-resolution spatial transcriptomics, and histology into a spatiotemporally unified representation that fuses molecular, functional, and structural information (Fig. 6a). Using this approach, we topographically map gene-expression states, local cellular composition, and circuit dynamics to reveal—across maturation stages—molecular determinants of neural output activity and collective oscillatory recruitment.

**Figure 6.**
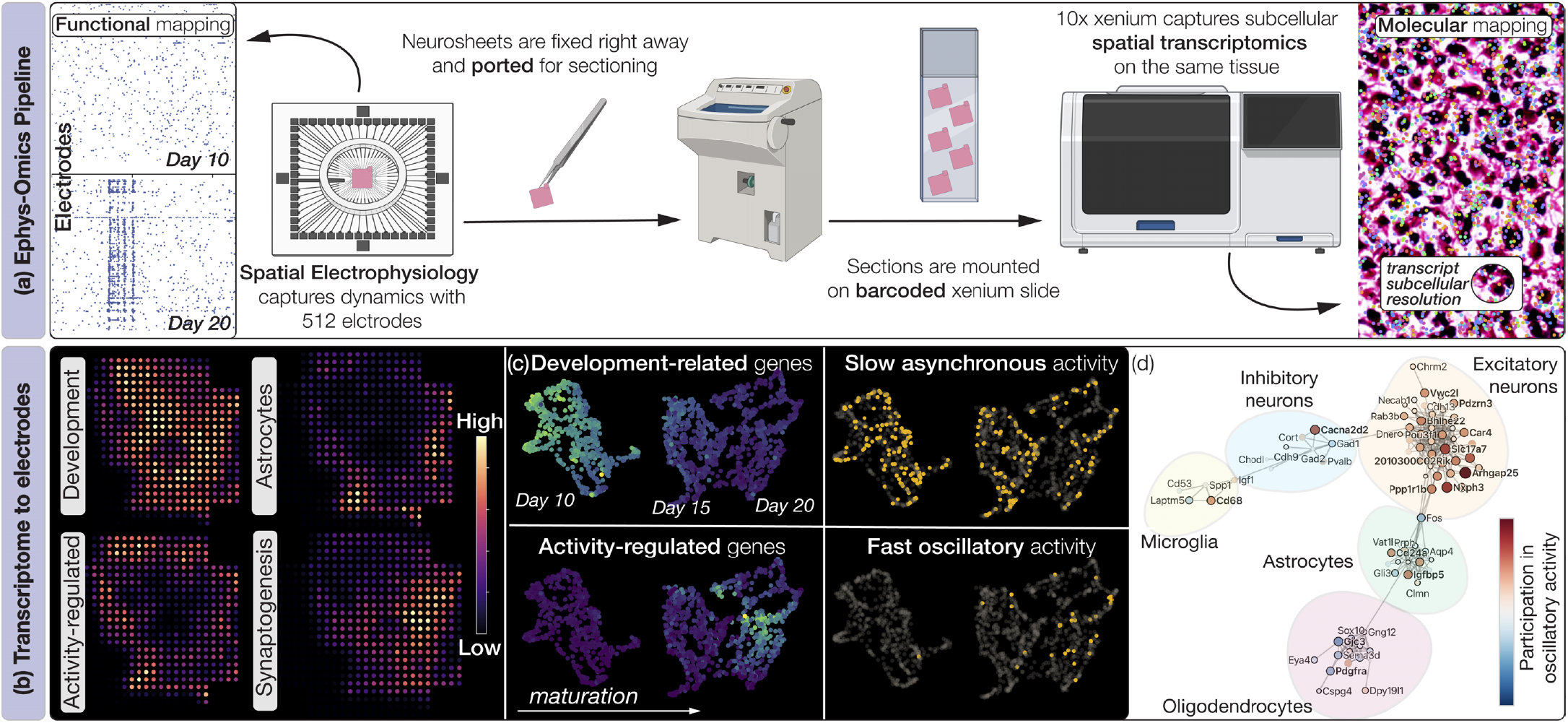
Joint electrophysiology and spatial transcriptomics. **(a)** The experimental pipeline involves performing high-resolution MEA electrophysiology on Neurosheets, followed by fixing and cryosectioning. The sections are harvested from the top to ensure capturing the cells that were previously in contact with the MEA and placed on Xenium’s barcoded slide. The slides are then processed and run through Xenium to obtain the spatial transcriptomics data.**(b)** Despite homogeneous cellular seeding, specific gene programs primarily occupy spatially distinct locations as shown by UMAP clusters. The clusters are shown on a representative Day 10 sample. **(c)** UMAP clustering further reveals age-specific gene clusters and systematic, temporal up- and down-regulation of activity- (Arc, BDNF) and development-related (Sox11, Nrep) genes, respectively. Dynamically, this results in a shift from slow asynchronous neural activity to fast collective oscillatory behavior represented here with their corresponding inter-spike intervals (Fig S15). **(d)** Gene-coexpression map revealed distinct cell-identity-specific clusters, colored by their spatial correlation with cellular density and, by proxy, with the network’s bursting/oscillatory activity (SI 8, Figs. S16-18).

Briefly, Neurosheets prepared with primary cortical neurons are aligned on MEAs and recorded to capture the evolution of spatiotemporal spiking dynamics across Days 10, 15, and 20 of maturation. Consistent with previous reports (*17, 35*), we observe a monotonic increase in firing rate as the tissue matures, together with a shift from predominantly slow asynchronous, local dynamics on Day 10 to fast oscillatory, globally bursting activity on Day 20 (Fig. 6a). Immediately after electrophysiological recording, tissues are fixed, cryoprotected, and sectioned to isolate the network layer from the apical surface, thus preserving access to cells in contact with the microelectrodes. Sections are then mounted on Xenium RNA-barcoded slides and processed for high-resolution spatial transcriptomic profiling of 248 brain-associated genes. After Xenium imaging, the same slides are processed for histology to provide structural information. Finally, transcriptomic data are mapped onto corresponding electrodes and subsequently projected onto histological images using alignment markers shared between tissue and MEA (Fig. 6b), resulting in co-registered, multimodal, multiscale measurements from the same tissue.

The mapped transcriptomic data reveal complex, nested, and spatially segregated genetic profiles within individual Neurosheets, despite their homogeneous preparation (Fig. 6c). These patterns point to the segregation of initially equivalent cellular populations into distinct transcriptional identities, a process likely governed by an intricate interplay among molecular machinery, cell-cell signaling, and activity-dependent feedback. Beyond spatial segregation within individual samples, key genetic programs also evolve in time. This temporal progression is illustrated in Fig. 6(c), which shows the joint embedding of electrophysiological-transcriptomic data obtained through UMAP clustering (Uniform Manifold Approximation and Projection (*53*), a dimensionality-reduction approach for representing complex multimodal data). The embedded programs exhibit a clear temporal shift over the timecourse of the experiment, gradually transitioning from development-related states toward activity-regulated states, thereby reflecting a functional progression in neural behavior from asynchronous activity to collective oscillatory dynamics.

As the tissue matures from Day 10 to Day 20, the analysis reveals a systematic downregulation of initially dominant development-associated programs, accompanied by progressive upregulation of activity-dependent programs. Because Neurosheets are generated from dissociated neural cells, the early predominance of development-related genes likely reflects a prioritization of network formation through neurite extension and synaptic scaffolding. This developmental phase may establish the structural substrate required for the emergence of collective neural dynamics, in turn inducing the upregulation of activity-based transcriptional programs, within a positive feedback loop. Overlaying single-electrode interspike-interval data from electrophysiological recordings onto the UMAP transcriptomic clusters points to this process (Fig S15(b)). Indeed, the data indicate that neural populations are recruited into fast spiking, synchronous oscillatory activity in regions associated with the upregulation of activity-dependent programs, after the downregulation of development programs.

Finally, gene-coexpression maps reveal coordinated organization among several gene clusters corresponding to stereotypical cellular classes, including excitatory and inhibitory neurons, astrocytes, microglia, and oligodendrocytes (Fig. 6(d)). While most genes remain associated with their respective clusters across time points, *Fos*, a canonical activity-regulated gene, initially localizes within astrocytic clusters and then progressively shifts to form a link between neuronal and astrocytic programs during later development, tracking the maturation of network dynamics (Fig. S15). Spatial gene-expression patterns also exhibit variable correlations with cellular density, with excitatory neuronal markers showing a positive association. Participation in spatial oscillatory activity or bursting likewise correlates positively with cellular density (Fig. S17)—and, by extension, with excitatory neuronal populations (Fig. S18). Inhibitory neuronal markers, by contrast, display a more heterogeneous pattern, with individual genes weakly correlated or anticorrelated with cellular density. This suggests a comparatively immature inhibitory population whose transcriptional programs have not yet become coherently coordinated, despite increasing coupling to excitatory neuronal programs (Fig. S15). Together, these findings highlight the role of excitation/inhibitory balance in coordinating oscillatory neural activity, and illustrate how such convergent analyses enabled by Neurosheet have the potential to shed new light on the complex multiscale processes at play.

## Discussion

Realizing living neural architectures that are scalable, robust, and functionally programmable requires biofabrication strategies that simultaneously control tissue realization, preservation, and interrogation. Neurosheets address this challenge by reimagining porous sheets—simple yet versatile materials—as programmable substrates for neural assembly. By leveraging the separation of scales between somas and neurites, sheets with pre-programmed porosity enable control over the spatial organization of cellular populations into complex architectures, obtained through cutting, buckling, folding, stacking, and modular composition. At the same time, they allow the emergence of 3D plastic networks typical of natural systems. This material-based and intuitive approach represents a conceptual advance in the design and study of neural circuit organization, geometry, and function, offering a reproducible platform in which emergent dynamics can be systematically explored and directed through deliberate tissue design. Coupled with custom bidirectional optoelectronics, microfluidic channels, biosensors, electronics, and mechanical actuators, Neurosheet technology establishes a versatile paradigm for harvesting neural dynamics at scale for scientific discovery, computation, and translational applications. Its portability further enables cross-platform convergent analyses, whereby electrical activity can be directly aligned with high-resolution spatial imaging and multiomic analysis, uniting molecular identity, structure, and function across scales. Neurosheets thus offer a reductionist, complementary alternative to low-throughput, poorly controllable, and often overly complex animal systems, while opening a path toward predictive, design-driven models of neural dynamics. With advances in multiomics, AI-enabled multimodal discovery, and scalable perturbation toolkits, this platform is positioned to close the loop between observation and design, allowing neural systems to be iteratively engineered, analyzed, and unraveled.

## Acknowledgements

This study was jointly funded by the NSF Expeditions in Computing ‘Mind in Vitro’ #2123781 (M.G.), NSF EFRI BEGIN OI #2515342 (M.G.), and the Scott H. Fisher Multi-Cellular Engineered Living Systems Theme at the Carl R. Woese Institute for Genomic Biology (M.G., T.S., H.K.). Spatial transcriptomics experiments were supported by the Grainger College of Engineering Strategic Research Initiative (S.M.). We thank Prof. Hee Jung Chung’s lab at UIUC for generously providing the primary rat hippocampal and cortical neurons. We thank Core Facilities at the Carl R. Woese Institute for Genomic Biology, Materials Research Laboratory Central Research Facilities, Holonyak Micro and Nanotechnology Lab, Beckman Institute, and Roy J. Carver Biotechnology Center at UIUC for technical assistance. Biorender was used to generate elements of Fig.6.

## Author contributions statement

G.U.and M.G. conceptualized and designed the research. G.U., K.K., X.Z., H.Y., Z.D., and S.K. performed experimental research. G.U., K.K., and S.H.K., L.M, A.D, and S.M. analyzed data. All wrote the paper. **Supplementary Information** is available for this paper. **Correspondence** should be addressed to M.G. **Correspondence** should be addressed to M.G.

## Data availability

The data supporting the results of this study are available in the main text, methods, and supplementary information. All hardware design files (MEA/PCB) are available at https://github.com/GazzolaLab/MiV-OH. All data are available upon request.

## Methods

### 1 Tissue fabrication and characterization

This section presents the biofabrication protocols and methods involved in the structural characterization of living neurosheets:

#### 1.1 Porous Sheets

In this work, we tested two commercially available porous sheets: cellulose acetate membrane filters (Sterlitech) and polycarbonate membrane filters (Sterlitech).

##### 1.1.1 Cellulose acetate membrane filters

As purchased, cellulose acetate membranes were already reinforced with supporting fibers mimicking a paper-like isotropic fibrous structure for structural support, Fig. S1(b). For cellulose acetate membranes, pore sizes of 0.8, 1.2, 3, and 5 *µ*m were tested.

##### 1.1.2 Polycarbonate membrane filters

Polycarbonate filters were alternatively selected due to their transparent nature, allowing brightfield visualization, a smaller thickness necessary for multilayer constructs, and non-fibrous cylindrical pores, Fig. S1(c). Here, a 20 *µ*m pore size was selected to minimize cell soma migration while allowing functional bidirectional connectivity through pan tissue neurite growth and interactions.

##### 1.1.3 Preparation

All sheets were obtained sterilely from the manufacturer and were subjected to oxygen plasma treatment before use, followed by three times washing with 1× phosphate-buffered saline (PBS) and air-drying under sterile conditions. The dried membranes were cut into defined geometries using a benchtop laser cutter (xTools M1). To minimize contamination, the laser cutter stage was sterilized with 70% isopropanol before cutting. The laser cutting parameters, such as power and speed, were optimized to minimize burning and maintain dimensional tolerance, Fig. S1(a). Topologies are then coated with 0.1 mg/ml Poly-D-lysine (Thermo Fisher Scientific) at room temperature for an hour to impart a positive surface charge and enhance adhesion of cells and hydrogels. Once coated, the samples are rinsed thrice with 1× phosphate-buffered saline (PBS) and air-dried before further use.

#### 1.2 Hydrogel preparation and integration with sheets

A mixed hydrogel composed of collagen type I and Matrigel was used in this study. As collagen and matrigel are thermoresponsive, all hydrogel manipulations were performed in an ice bath and with chilled components (pipette tips, vials, buffers). Briefly, Collagen type I, rat tail (Corning) was neutralized to pH 7.2–7.4 with NaOH buffer and adjusted to a final concentration of 4 mg/mL. The collagen solution was then combined with high concentration, growth factors reduced Matrigel (Corning, 10 mg/mL), and neural maintenance medium (NMM) to obtain the desired final concentrations. Unless specified otherwise, experiments were carried out with a 3 mg/mL Collagen : 1 mg/mL Matrigel formulation (total 4 mg/mL protein). Since neurite outgrowth was found to be maximized at 2 mg/mL total hydrogel concentration, Fig. S1(g), a 3:1 collagen-to-Matrigel ratio at 2 mg/mL was used for bilayer and multilayer cases. Use of pure matrigel as hydrogel at 4mg/ml was also tested but resulted in excessive cellular clumping, Fig. S1(e), hence it was not pursued.

Once the hydrogel is ready at the desired composition, it is poured onto the porous sheets, which in turn are placed atop a cured PDMS surface to create a hydrophobic-hydrophilic wettability contrast. Following coating, the hydrogel was allowed to partially polymerize under high-humidity conditions (maintained by placing water droplets near the samples) at 37 °C for 10 min.

#### 1.3 Cell culture

In this study, we tested neurons from a range of sources to demonstrate the robustness of our approach. The key cell types tested are described below.

##### 1.3.1 Embryonic stem cells derived neurons

We tested two mouse embryonic stem cell lines, wild-type AB2.2, and optogenetic ChR2-HBG3 Hb9-GFP. The differentiation protocols remain constant for both cell lines.

**Motoneurons** were derived according to protocol highlighted in (*54*). Briefly, mouse embryonic stem cells (mESCs) were defrosted and expanded in a feeder culture with mouse embryonic fibroblasts (MEFs). Media was exchanged every 24 hours with fresh Leukemia inhibition factor (ESGRO mLIF, Sigma Aldrich) to prevent unwanted differentiation. Once the colonies are large enough, they are detached with 0.05% Trypsin-EDTA and dissociated to a single-cell suspension. Trypsin is neutralized with serum-containing medium, and cells are centrifuged. The cells are resuspended in MEF growth media and plated in a T-175 flask (Fisher Scientific) to separate MEFs from mESCs. Approximately 45 mins-1 hour later, floating cells are collected, centrifuged, and resuspended in Neural induction media (NIM) at 2M cells per 100 mm dish for two days. The cells reaggregate as embryoid bodies during this duration. From Days 2-6, the embryoid bodies (EBs) are cultured in Neural Differentiation Media (NDM). On day 7, the EBs are collected and gently dissociated using Accumax (ThermoFisher) to a single cell suspension. The cells are then plated in NDM supplemented with 20% Horse Serum for two hours to filter out non-adherent cells. The attached cells are collected with Accutase (ThermoFisher) and plated on sheets as needed.

##### 1.3.2 Primary neurons

Fresh primary cells were obtained from Dr. Hee Jung Chung’s lab at UIUC. Briefly, hippocampal and cortical areas were dissected from time-pregnant rats at E18-E19, and put in MilliQ water (pH 7.4, 4 °C) supplemented with 1.16% Na2 SO4, 0.52% K2 SO4, 0.24% 4-(2-hydroxyethyl)-1-piperazineethanesulfonic acid (HEPES), 0.18% d-glucose, and 0.1% MgCl2 6H2O. Neurons were further dissociated and kept in Minimum Essential Medium (MEM) Eagle’s with Earle’s Balanced Salt Solution (BSS) without l-glutamine, supplemented with glucose, 100 mM sodium pyruvate, 200 mM l-glutamine, and 100 U/ml penicillin and streptomycin. As an alternate source, perhaps more amenable to general labs, primary mouse cortical neurons were directly procured from thermofisher (Catalog No. A15586), defrosted, and seeded on sheets following protocols previously described for the other cell types.

#### 1.4 Media Formulations

##### 1.4.1 MEF Growth Media

MEFs were plated a day before the seeding of stem cells. DMEM (high glucose, no l-glutamine) serves as the basal media, supplemented with 10% fetal bovine serum, Glutamax, non-essential amino acids, and Pen/Strep.

##### 1.4.2 Stem Cells Growth Media

Mouse embryonic stem cells were grown atop MEF feeder layer. The expansion media consists of Embryomax DMEM as basal media, supplemented with 10% embryonic stem cell certified fetal bovine serum (FBS), Glutamax, Pen/Strep, Nonessential amino acids, and Embryomax nucleosides. The media was exchanged every 24 hours and freshly supplemented with mouse LIF and 2-mercaptoethanol.

##### 1.4.3 Neural Induction Media

Dissociated stem cells are resuspended in neural induction media to form cellular aggregates/embryoid bodies and impart neural lineage during the first two days of differentiation. The media is 1:1 Advanced DMEM/F12: Neurobasal media supplemented with 10% Knockout replacement serum, Glutamax, and Pen/Strep. The media is freshly supplemented with bFGF (20ng/ml), Noggin (20ng/ml), FGF-8 (20ng/ml), and 2-mercaptoethanol.

##### 1.4.4 Neural Differentiation Media

For the next 7 days after neural induction, the embryoid bodies were cultured in Neural Differentiation media with fresh media replacement every two days. The media is 1:1 Advanced DMEM/F12: Neurobasal media supplemented with 10% Knockout replacement serum, Glutamax, and Pen/Strep, B27 supplement, N2 supplement, ITS supplement, and heparin. The media is freshly supplemented with Retinoic acid (1*µ*M), Ascorbic acid (100 *µ*M), smoothened agonist (1*µ* M), and 2-mercaptoethanol.

##### 1.4.5 Neural Maintenance Media

Once fabricated, all the tissues were maintained in Neural Maintenance Media (NMM) comprising neurobasal plus media supplemented with B27 plus supplement, Pen/Strep, and Glutamax. For the first three days of tissue culture. The media is supplemented with a growth factor cocktail consisting of BDNF, GDNF, NT3, CNTF, BFGF (10ng/ml each), Forskolin and IBMX.

#### 1.5 Culturing cells on sheets

Once the sheets are prepared and infused with hydrogel, the cells are seeded in the form of a suspension from the top. The sheets are kept on hydrophobic PDMS, which ensures the liquid beads up around the sheet area, ensuring robust control over the cell-seeded area. The volume of suspension must be calibrated to obtain the desired cell seeding density and ensure all of the sheets’ area is properly wetted. Too high a volume can further lead to the leaking of cell suspension outside the sheet’s surface onto the PDMS. This procedure is kept constant for all the tissue cases and cell types used. To promote initial attachment, the cells are seeded in NDM supplemented with 20% horse serum and are allowed to adhere to the surface for 2-3 hours. Once attached, the tissues are inundated with Neural Maintenance Media supplemented with necessary growth factors, and the tissues are allowed to mature.

In the case of the out-of-plane tissues, the tissues are folded before hydrogel introduction. To accommodate the out-of-plane regions, for instance, the buckled region of the arch bridge, holes were punched in the PDMS. This ensures legs of the tissue are flattened and make perfect contact with the PDMS layer for cell seeding. The cells are next seeded from the top and allowed to attach for 2-3 hours.

In the case of bilayer and multilayer constructs, the sheets are sandwiched between layers of through-punched PDMS to ensure easy flipping and no leakage of cells from the sides. The hydrogel is imbued from both sides to ensure full coverage, and the cells are seeded one side at a time. Once the cells on one side of the tissues are attached, the PDMS construct is flipped, and seeding is performed on the other side. For multilayer constructs, multiple layers are placed atop each other with fresh collagen-matrigel (3:1, 2mg/ml) mix in between and sandwiched between the layers of PDMS, similar to ones used for forming the bilayer tissues. After 30 minutes of gelation, the tissues are inundated with neural maintenance media and cultured as previously described.

#### 1.6 Viability

To assess the viability of neurosheets, a commercially available kit (LIVE/DEAD Viability/Cytotoxicity Assay Kit (Green/Deep Red), ThermoFisher) was used according to the manufacturer’s protocol. Briefly, live cells are labelled using cell-permeant non-fluorescent Calcein that becomes intensely fluorescent calcein with the intracellular esterase activity, producing an intense uniform green fluorescence in live cells (ex/em: 495/515 nm) detectable with GFP filter set. To label dead cells, SYTOX deep red is used, which enters the cells with damaged membranes and exhibits increased fluorescence on binding to nuclei, which can be detected with a CY5/Deep Red filter set (ex/em maxima: 660/682 nm).

The samples were collected on day 14 of maturation and incubated with the dyes at a 1:2000 dilution in maintenance media for 30 minutes. The cells were washed with PBS and imaged right away. The microscopy was performed on a Zeiss confocal laser scanning microscope, LSM 900, using a 20X objective. Most of the samples routinely exhibited viability higher than 85% irrespective of the initial seeding density for stem cell-derived cultures, Fig S1(f). The minimal loss in viability can be attributed to induced stress during the spheroid or tissue dissociation process.

#### 1.7 Second harmonic generation

To characterize the extent of hydrogel inundation inside the tissue and collagen microstructure post-gelation, we used nonlinear second harmonic generation microscopy. The hydrogel-infused sheets were prepared as previously described and were mounted on the Zeiss confocal laser scanning microscope, LSM 710, coupled with a Ti-Sapphire Laser (Spectra Physics Mai Tai) 690-1040 nm and SHG filter set to perform imaging. Fig S1(d) shows a representative image where the reinforcing fibers from the cellulose acetate membrane filter can be seen alongside collagen type 1 fibers creating a dual scale architecture, necessary for both micro-and meso scale control.

#### 1.8 Scanning electron microscopy

While fluorescence imaging provides high-resolution cellular visualization, it does not fully capture the microarchitecture of the hydrogel matrix or sheets. To enable simultaneous visualization of both neural tissue and its structural environment, scanning electron microscopy (SEM) was performed using a Quanta FEG 450 ESEM (FEI). On day 14 of maturation, tissues were gently rinsed in 1× PBS and fixed in 2.5% glutaraldehyde and 4% paraformaldehyde in 0.1 M sodium cacodylate buffer for 4 h at 4 °C. Following fixation, samples were washed three times in PBS (5 min each) and dehydrated through a graded ethanol series. Samples were dried using an Autosamdri-931 critical-point dryer (Tousimis), mounted onto aluminum SEM stubs using carbon adhesive tape, and sputter-coated with a 10 nm gold layer. Imaging was performed immediately after preparation. SEM was used to structurally characterize the neurosheets, highlighting the porous sheet’s architectures, hydrogel, neural cells, and their processes on both the cell-side and the opposite side, as shown in, Fig.1d.

#### 1.9 Immunostaining

To perform immunostaining, tissues were collected and washed three times (5 mins each) with 1X Phosphate Buffer Saline(PBS). Next, the samples were fixed by immersing them in 4% freshly prepared Paraformaldehyde (PFA) at room temperature for 20 minutes. Once fixed, excess PFA is removed, and tissues are subsequently washed three times as before. Permeabilization of cellular structures is achieved by immersing tissues with 0.1% Triton-X for 5-10 minutes, followed by washing and a one-hour incubation of tissues in 2% Bovine Serum Albumin (BSA) blocking buffer at room temperature. Post-washing the buffer solution, primary antibodies for detecting neurons (anti beta 3 tubulin, rabbit, Abcam, 1:500) and astrocytes (Anti Glial Fibrillary acidic protein, mouse, Millipore, 1:2000) are diluted in 0.1% BSA and incubated overnight at 4 °C. Next, secondary antibodies from the Alexa Fluor series are utilized to bind fluorescent labels to primary antibodies. All manipulations for secondary antibodies are performed in the dark. The antibodies are diluted in 0.1% BSA at 1:1000 and applied to the tissues for two hours at room temperature, followed by washing and nuclear staining with DAPI for 10 minutes. At this point, samples can be directly imaged; however, refractive index matching can further improve the depth of tissue that can be imaged. Such clearing is critical to imaging multilayer constructs with confocal microscopy and is achieved by immersing tissues in RapiClear 1.49, a water-soluble tissue clearing medium.

Immunostaining was performed across tissue types and batches to ensure homogenous distribution of neurons and astrocytes on the tissue surface and neurites throughout volume, Fig.1,3,4,5., and Fig. S2, S5

#### 1.10 Cryosectioning

Tissues were fixed as described above and sequentially incubated in 5%, 10%, 15%, 20%, 25%, and 30% (wt/vol) sucrose diluted in 1× PBS for 6 h each at 4 °C. Samples were incubated overnight in 30% sucrose to ensure complete infiltration. Cryoprotected tissues were embedded in optimal cutting temperature (OCT) compound (Tissue-Tek) in cryomolds and snap-frozen. Sections were prepared on a cryostat (Leica CM3050 S) and collected onto Superfrost™ Gold slides (Thermo Fisher Scientific). Slides were allowed to air-dry for 1 h at room temperature before subsequent immunostaining or histological staining. Cryosectioning was used to visualize the extent of neurites within the neurosheets’ volume for bilayer tissues prepared with cellulose acetate membrane and wild-type mESCs-derived motoneurons as shown in Fig. S2

### 2 Electrophysiology Methods

In this section, we describe the necessary protocols and analysis details for performing both imaging-based and microelectrode arrays (MEA) based electrophysiology on living neurosheets:

#### 2.1 MEA fabrication

All the MEA layouts utilized throughout this study are visualized in Fig. S3 and were designed using KLayout and are open-sourced here (*55*). The microelectrodes were fabricated as described in (*17*). Briefly, we select 3-inch borosilicated glass wafers, which were spin-coated with AZ5214E photoresist (PR) using a Headway PWM32 spin coater to get a 1 *µ*M thick coating. Maskless photolithography was performed using Heidelberg MLA 150 Maskless Aligner through a 375 nm laser source with a dose of 210 mJ cm^*™*2^. The pattern was then developed using 1:4 diluted AZ400K developer, washed with water, and dried by blowing air before baking for 50s at 120 °C. Next, dual-layer metal deposition was performed with 15nm Titanium and 85 nm Platinum using the AJA sputter coater. Post-metal deposition, liftoff was performed using acetone, followed by silicon nitride passivation layer deposition using Oxford PECVD.

The second lithography was performed the same way as the first one, with spin coating the PR, pattern exposure, developing, and baking to selectively etch the passivation layer at the electrodes and contact pads using CCF4-based Reactive Ion Etching (Plasmatherm 790 MF). Post RIE, the MEAs are washed with acetone to remove excess photoresist and tested for impedance to ensure a high signal-to-noise ratio. Typical electrode impedances for the MEAs used in this study ranged from 500 kΩ to 1.5 MΩ, measured at 1 kHz.

#### 2.2 Data collection

Extracellular recordings were acquired using the Mind in Vitro electrophysiology platform (*17*). The acquisition pipeline consists of custom-fabricated printed circuit boards (PCBs) interfaced with the MEA through spring-loaded POGO-pin contacts, enabling direct electrical coupling to the tissue-mounted microelectrodes. Signals were routed to the OpenEphys acquisition box through serial peripheral interface (SPI) cables coupled to Intan headstages and integrated amplifier chips, which provided high-impedance buffering and low-noise amplification. OpenEphys GUI was used to acquire the raw data at 30KHz without online filtering to preserve raw statistics for offline analysis. Before data acquisition, electrode impedances were measured to verify reliable contact between the POGO-pin interface and the MEA contact pads, and to identify and exclude high-impedance or non-functional channels.

#### 2.3 Optical Stimulation

Optical stimulation was performed using optical fibers (Doric; Mono Fiber-optic Patch Cords, MFP 480/500/-0.63, 1 m, FC-MF1.25(F)) mounted upright and coupled to laser diodes (Doric). Stimulation parameters, including stimulus period, frequency, number of pulses, and total stimulus duration, were programmed using Doric Neuroscience Studio. For Y tissues, in which concurrent stimulation was delivered, the two stimulation patterns were programmed separately, with the onset of the trailing pattern triggered via TTL-based coupling to ensure high temporal fidelity.

#### 2.4 Electrical Stimulation

Electrical stimulation was delivered using an Intan RHS Stim/Recording System. The system was interfaced with the Mind in vitro platform’s 128-electrode PCB via four RHS 32-channel Stim/Recording Headstages. For sandwich MEA experiments, a 64-electrode PCB was used with two headstages. Data acquisition and stimulation patterns were programmed using Intan RHX software. Stimulation consisted of biphasic pulses of 1–2 ms duration, with stimulus amplitude adjusted between 500 nA and 2 *µ*A to elicit robust multichannel neural responses.

#### 2.5 Chemical Stimulation

Chemical stimulation was assessed by application of the AMPA receptor antagonist DNQX (6,7-dinitroquinoxaline-2,3-dione) at 10 *µ*M concentration diluted in NMM to wild-type motoneuron-derived neurosheets. Neural activity before and after DNQX application was compared and showed a marked reduction in periodicity, Fig. S6(b).

#### 2.6 Data Analysis Pipeline

The electrophysiological recordings were obtained using the Open Ephys acquisition terminal and subsequently processed using MiV-OS, an open source Python platform developed by the Gazzola lab for neural data analysis (*56*). Computational post-processing and batch analysis were performed using allocated resources on the Frontera supercomputer. The raw data were first loaded into the analysis pipeline and then bandpass-filtered using a fourth-order Butterworth filter with cutoff frequencies set at 300 Hz and 3000 Hz, filtering out slow field potentials and high-frequency noise for a cleaner spiking signal. Spike detection was then performed by applying a threshold at 5median{x}/0.6745, where x denotes the bandpass-filtered signal. Raster plots were employed to visualize both spontaneous and stimulus-evoked neural activity based on detected spike times, with the y-axis representing the individual recording channel for each customized MEA design.

##### 2.6.1 Spatial heatmaps

To visualize spatial patterns of neuronal firing across the tissue, each channel/electrode’s spike train was binned into 25 ms intervals, and spike counts were measured for every recording electrode to compute the local firing rate within each time window. The resulting firing rates were further interpolated using a Gaussian kernel (with *σ* = 0.16*mm*) and projected onto the layout of the designed MEA to generate continuous spatial heatmaps, providing a two-dimensional representation of network activity. The colormap was power-normalized with gamma = 0.7 to better highlight the activity propagation and underlying dynamical structures. Together, these heatmaps allow us to assess the spatiotemporal dynamics both during spontaneous activity and stimulus-evoked dynamics.

##### 2.6.2 Peristimulus time histograms (PSTH)

Peristimulus time histograms (PSTHs) were generated to quantify the temporal dynamics of the network’s response to optical and electrical stimulation. Spike trains were aligned to the onset of stimulation pulses detected via TTL signals and binned into fixed-width time bins (2 ms unless specified otherwise). For each electrode, spike counts within each bin were averaged across repeated stimulation trials and normalized by bin width to obtain mean firing rates. Population PSTHs were then computed by averaging firing rates across predefined spatial groups of electrodes corresponding to distinct regions of the neural tissue. For comparing the same sample performance on multiple days, as shown in Fig.2, maintext, Normalized PSTHs are plotted by dividing the PSTH values by mean firing rates from the spontaneous data.

#### 2.7 Traveling wave characterization-Ring Tissue

The ring tissues were optically stimulated using a 4-pulse and an 8-pulse protocol at 40 Hz with a 3-second interval between each stimulation bout, Fig. S4. For each stimulation trial, spike trains from all 480 channels were binned into 25-ms intervals to compute firing rates over a 1-s post-stimulus window. The maximum firing time after stimulation for each channel was then identified, representing the moment of peak local activation. To exclude electrodes with negligible activity, only electrodes whose peak firing rate exceeded 1.2 times their mean firing rate over the analysis window were considered. These peak times were averaged across each radial row to obtain a single mean activation latency per row, effectively capturing how each radial segment of the tissue responded to stimulation.

This procedure was repeated across all stimulation instances and locations, and the resulting row-wise timing values were overlaid to visualize propagation dynamics across the ring geometry. The displayed points in Fig. S4(b,c) correspond to these row-averaged peak times, with color shading (green for 4 pulse and purple for 8 pulse trials) indicating the order of stimulation instances (darker shades representing later trials). The red dots mark the mean activation time across all trials for each row, and linear fits were applied to these averages to estimate the slope of activity propagation across the tissue, capturing the direction and speed of propagation within the tissue (*≈*57 mm/s).

#### 2.8 Calcium imaging

Intracellular calcium activity was monitored using Calbryte™ 520 AM (AAT Bioquest), a green fluorescent calcium indicator. Tissues were gently removed from culture dishes and incubated in 5 *µ*M dye diluted in neuronal maintenance medium (NMM) for 45 min at 37 °C. Following incubation, samples were rinsed and immersed in fresh pre-warmed NMM, and imaging was performed. Planar cultures were imaged on an epifluorescence microscope (Nikon Eclipse Ti) equipped with a fluorescein isothiocyanate filter set, with an excitation/emission wavelength of 490/525 nm. Recording was conducted under a 488 nm laser illumination, acquiring 10 frames/s for 1 min. For thicker constructs (bilayer and multilayer tissues), calcium imaging was performed using a confocal microscope, LSM 900, Zeiss, to enable volumetric sampling and minimize scattering-related signal loss, SI Video 12.

Alternatively, the Zeiss lightfield 4D microscope was utilized to simultaneously image calcium activity in multiple layers to assess the multilayer functional connectivity, SI Video 13.

#### 2.9 Imaris

Imaris software (version 10.1.1; Bitplane) was used to generate 3D animation videos of ring-shaped and Y-shaped tissues. The software was further used for filament reconstruction of ChR2-Hb9-GFP–labeled motoneurons derived multilayer tissues. Filament reconstructions were performed using the Imaris Filament Reconstruction module. Briefly, the Tree Autopath algorithm was applied to detect neuronal filaments, with an estimated soma diameter of 10 *µ*m. Intensity thresholding parameters were optimized to ensure reliable detection of neuron somas across all three imaged tissue layers while minimizing background signal. Next, automated filament tracing was performed with a minimum filament diameter of 1 *µ*m. AI-assisted filament detection within Imaris was subsequently used to identify and exclude spurious filament reconstructions; however, a few artifacts may remain. Final filament renderings were generated and colored based on Z-height statistics.

Additionally, Imaris was used to render Z-stack fluorescence images of the arch-bridge and cross-bridge tissues. During imaging, tissues were gently compressed under a coverslip, resulting in a reduced imaged Z-height relative to the original tissue height. Accordingly, the acquired Z-stack depths were 343 *µ*m for the arch-bridge sample and 174 *µ*m for the cross-bridge sample.

The entire stack was imaged with 10 equally dispersed Z locations. For visualization purposes only and to better highlight the folded regions, the cross-bridge dataset was scaled linearly in the Z-dimension to a final depth of 800 *µ*m in Imaris.

### 3 Matching Spatial Tissue Transcriptome with Spatiotemporal Electrical Activity

This section describes extended details on our experimental and analytical approach involved in generating the same tissue electrophysiology, spatial transcriptomics, and histology dataset using Neurosheets’ cross-platform portability.

**Tissue Preparation:** Square neurosheets (4 mm*×* 4 mm) with an extended orientation marker were prepared from primary rat cortical neurons. Each neurosheet contained 150 K cells and was fabricated using the procedure described in Section 1.5.

#### 3.1 Electrophysiology

Neurosheets were matured for 10, 15, and 20 days in Neural Maintenance Medium. On the day of the experiment, the MEA recording device and the surrounding environment were cleaned with RNase Away (ThermoFisher) to avoid RNA/DNA contamination. Neurosheets were placed on the MEA, oriented using the extended marker, and aligned under a microscope to ensure the sheet was centered on the electrode area. To minimize contamination risk, Kimwipes were not used to press down the tissue. Tissue activity was allowed to stabilize for 5 minutes and was then recorded for 2 minutes. Tissues were carefully unmounted after electrophysiology, washed with PBS (3*×*), and fixed in 4% PFA at room temperature for 30 minutes. This procedure was repeated for 16 samples per day, totalling 48 samples for the entire experiment across 3 time periods.

##### 3.1.1 Cryosectioning

After fixation, tissues were washed and sequentially incubated in 5, 10, 15, 20, and 30% sucrose solutions for 2 hours each. After exchanging the final 30% solution with fresh 30%, samples were kept overnight at 4 °. The next day, samples were placed in cryomolds (Tissue-Tek) with Tissue-Tek^®^ O.C.T. Compound and snap-frozen. OCT-embedded tissues were mounted on a cryostat and sectioned into 10 *µ*m thick slices. The initial tissue sections, comprising the network layer directly in contact with MEAs, were collected and mounted on the Xenium Slide according to the orientation marker. Subsequent tissue sections from the same tissue were assessed for RNA quality by DV200.Xenium Run: Cryosectioned slices were processed for Xenium spatial transcriptomics following 10x Genomics protocols adapted for fixed-frozen input, drawing on the Visium CytAssist fixed-frozen tissue-preparation and rehydration guides (CG000663, CG000662), the Xenium FFPE decrosslinking guide (CG000580), and the standard Xenium assay and analyzer workflows (CG000582, CG000584). All buffers (1X PBS, PBS-T) were prepared fresh, and Xenium FFPE Tissue Enhancer was thawed (30 min, 37 °, 300 rpm) before retrieving slides. Slides were rapidly thawed from dry ice on a 37 ° thermocycler adaptor (1 min), then rehydrated through a graded series at room temperature (1X PBS, 5 min; Milli-Q water, 3 min; 100% then 70% ethanol, 3 min each; Milli-Q water, 20 s), assembled into Xenium cassettes, and held in PBS-T. Tissue was decrosslinked at 22 ° using Decrosslinking Buffer (Xenium FFPE Tissue Enhancer with Diluted Perm Enzyme B), followed by three PBS-T washes (1 min each). Slides then proceeded immediately to probe hybridization, ligation, and amplification, and were decoded and imaged on the Xenium Analyzer without modification to the standard workflow. A pre-designed Xenium Mouse Brain Gene Expression panel (248 genes) was used. Cross-reactivity with rat was assessed using a mouse–rat cross-species hybridization efficiency report provided by 10x Genomics, reporting the fraction of probe sets per gene predicted to bind the rat ortholog. Of 248 targets, 188 (75.8%) were predicted to bind the rat ortholog fully and 216 (87.1%) at *≥*0.85 (mean 0.91). Due to the limited area on Xenium slides, we were able to mount 11 Neurosheets sections corresponding to various days, with a loss of one section during processing. The xenium run hence completed with 10 samples. Exact reagent volumes, catalog numbers, and incubation programs are as specified in the corresponding 10x Genomics protocol booklets.

#### 3.2 Post Xenium H&E staining

Following the Xenium Analyzer run, slides were processed for Hematoxylin & Eosin staining per the Post-Xenium Analyzer H&E Staining protocol (CG000613), right after the run. Autofluorescence quencher was first removed by immersing slides in freshly prepared Quencher Removal Solution (10 mM sodium hydrosulfite, used within 10 min) for 10 min, followed by three Milli-Q water washes (1 min each), all in a fume hood. Slides were then stained through a graded series at room temperature: Milli-Q water (2 min), Mayer’s Hematoxylin (20 min), three Milli-Q water rinses (1 min each), Bluing Solution (1 min), Milli-Q water (1 min), 70% then 95% ethanol (3 min each), Eosin (2 min), 95% ethanol (2 × 30 s), 100% ethanol (2 × 30 s), and xylene (2 × 3 min). Stained slides were coverslipped with 150–200 *µ*l mounting media, dried for 30 min at room temperature, and imaged. Exact reagent volumes, catalog numbers, and incubation details are as specified in the corresponding 10x Genomics protocol booklet (CG000613).

#### 3.3 Activity metric selection

For the primary electrophysiological metric, we chose network burst firing rate(which we will refer to as burst firing rate), as opposed to metrics like mean firing rate and non-burst firing rate. This is because burst firing rate(FR) was found to be strongly reproducible across bursts, making it a favorable metric, Fig S12.

#### 3.4 Transcript quality filtering

All downstream analyses were restricted to high-quality transcripts, defined as those with a Quality Value (QV) above 20. At the cell level, only cells with 15 or more qualifying transcripts were retained. At the electrode level, only electrodes with at least 100 such cells within a 150 *µ*m radius were carried forward, giving us an effective minimum transcript count of 1,500. This transcript count threshold was determined empirically.

#### 3.5 Generating pseudo-bulk expression profiles and UMAP

The Xenium spatial transcriptomics data exhibited considerable noise at the single-cell level, enough to prevent reliable cell type annotation, as shown in Fig S14. We therefore adopted a pseudo-bulk strategy, aggregating all cells within a 150 *µ*m radius of each electrode into a single transcriptional unit and normalizing expression by the total transcript count within that neighborhood.

#### 3.6 Generating the co-expression graph

The co-expression matrix was computed across all cells with at least one high-quality transcript (QV *>* 20). Pairwise co-expression was quantified using the Pearson correlation coefficient of raw transcript counts; edges were drawn between gene pairs with a coefficient of ≥ 0.45. Genes with zero variance across the dataset were excluded before computation. Alternative expression representations, including size-factor normalized counts and log-transformed expression, were evaluated and yielded qualitatively similar results. Nodes were colored by their log–log density-expression fits as shown in Fig S16.

#### 3.7 Activity vs. Density and Gene Expression vs. Density Plots

To characterize the relationship between network activity and spatial transcriptomics, we utilized local cellular density as a primary proxy, analyzing its correlation with both electrode activity and gene expression levels. We first identified network bursts by dividing the 120-second observation period into 0.02-second bins and selecting intervals where the total spike count across all electrodes exceeded 100. Because electrode activity during these bursts varied from sample to sample, we calculated a normalized firing rate for each electrode relative to the most active channel. To integrate this electrical data with spatial features, we defined local cell density by counting the number of cells within a 150 *µ*m radius of each electrode, restricting our analysis to cells with non-zero transcript counts in the spatial transcriptomics data. Within this same 150 *µ*m radius, we calculated fractional gene expression by normalizing the count of high-quality transcripts (Quality Value or QV≥20) for specific genes against the total transcript count across all genes in the region. Finally, electrodes were filtered to retain only those that exhibited a non-zero normalized firing rate during network bursts and possessed at least 50 cells and at least 1500 transcripts in their local neighborhoods. Finally, we plotted both the normalized firing rate during network bursts against local cell density (Fig S17) and the fractional gene expression against local cell density (Fig S18).

